# Computational generation of long-range axonal morphologies

**DOI:** 10.1101/2024.10.16.618695

**Authors:** Adrien Berchet, Remy Petkantchin, Henry Markram, Lida Kanari

## Abstract

Long-range axons are fundamental to brain connectivity and functional organization, enabling communication between different regions of the brain. Recent advances in experimental techniques have yielded a substantial number of whole-brain axonal reconstructions. While most previous computational generative models of neurons have predominantly focused on dendrites, generating realistic axonal morphologies is challenging due to their distinct targeting. In this study, we present a novel algorithm for axon synthesis that combines algebraic topology with the Steiner tree algorithm, an extension of the minimum spanning tree, to generate both the local and long-range compartments of axons. We demonstrate that our computationally generated axons closely replicate experimental data in terms of their morphological properties. This approach enables the generation of biologically accurate long-range axons that span large distances and connect multiple brain regions, advancing the digital reconstruction of the brain. Ultimately, our approach opens up new possibilities for large-scale in-silico simulations, advancing research into brain function and disorders.

## 1 Introduction

The brain is a highly complex organ responsible for controlling essential functions such as sensory processing, cognition, memory, respiration, motor control, and language production (Ito et al., 2020; Adamovich et al., 2024; Iranmanesh et al., 2021). These functions rely on intricate computations within distinct brain regions and the transmission of signals between them. Long-range axons, also referred to as projecting axons, control this inter-regional communication (Liu et al., 2024; Shin et al., 2020) and numerous studies have highlighted their critical role in the global neural circuitry (Zingg et al., 2014; Hilgetag and Zikopoulos, 2022; Mateus et al., 2024). Therefore, understanding the organization and connectivity of long-range axons is vital for deciphering brain function.

Recent advances in experimental imaging techniques have enabled the reconstruction of complete long-range axonal morphologies (Economo et al., 2019; Winnubst et al., 2019). However, the process remains time-consuming and costly, making it impractical to capture large portions of the biological neural circuits (Economo et al., 2019; Wang et al., 2018). In addition, experimental approaches face limitations in manipulating neuronal morphological properties to assess their effects on brain function. To overcome these challenges, numerical simulations of large-scale neuronal circuits are essential for gaining deeper insights into brain organization and the distribution of tasks across different brain regions. These simulations should model a large number of neuronal morphologies, including both dendrites and axons, to establish complete circuit connectivity. A major challenge is generating realistic neural circuits in which the modeled morphologies exhibit local morphometric properties (e.g., branching patterns and section characteristics) that closely resemble those of experimentally reconstructed neurons.

Long-range axons must target specific brain regions and follow pre-defined pathways, depending on the location of their source and targets (Brodal, 2010; Osten and Margrie, 2013). Various models have been proposed to synthesize artificial neuronal morphologies, each offering different tradeoffs. Some models focus on simulating a more realistic axonal growth during brain development, incorporating detailed molecular cues to guide the growth cone (Zubler and Douglas, 2009; Torben-Nielsen and Cuntz, 2014). While these models can reproduce detailed morphologies, they come with significant computational costs and require extensive data, which is difficult to acquire for large-scale circuits. On the other hand, mathematical models generate morphologies using mathematical principles and morphological statistics. Although these models require less experimental data, they need manual parameter adjustments based on the type and location of the neurons being synthesized, therefore harder to generalize to our use-case of axonal shapes (Koene et al., 2009; Luczak, 2006; Ascoli et al., 2001; Cuntz et al., 2010).

A recent model successfully integrated the mathematical and statistical approaches into a topological synthesis algorithm (Kanari et al., 2022). However, this work was limited to the synthesis of dendritic morphologies, due to the distinct nature of dendrites and axons. Local axons extend within a brain region and similarly to the dendrites they can be simulated taking into account local contextual cues. Long-range axons extend over large distances from the soma and target specific brain regions during development to form functional connections (Dickson, 2002; Sakai and Kaprielian, 2012; Kerstjens et al., 2022; Goodman and Shatz, 1993). Long-range axons must grow taking into account the distance from the soma, as dendrites and local axons do, but also navigate to specific target regions within the multiple brain regions (Kollins et al., 2009; Liu et al., 2024; Winter et al., 2023; Zhou et al., 2023). This requirement, along with the need for computational efficiency and realistic local morphometrics, cannot be fully captured by previous models.

In this work, we introduce a novel method for computationally synthesizing longrange axons that reproduce local morphometrics statistically similar to those of reconstructed axons while also targeting the appropriate brain regions based on the soma’s location. Our approach utilizes the Steiner Tree Algorithm to generate a skeleton connecting large-scale target regions, followed by a random walk guided by this skeleton to synthesize the primary axonal trunk. Finally, a local-scale model, based on topological synthesis (Kanari et al., 2022) is applied to generate the terminal branches of the axonal tree. We demonstrate that this method successfully combines large-scale target precision with detailed local-scale axonal synthesis, ensuring accurate targeting of brain regions while preserving realistic local morphometrics.

The organization of the paper is as follows. In Section 2, we briefly define the main concepts used in the following sections, then we describe the method in Section 3, and finally, in Section 4 we present several results to validate the method and propose an application in a mouse brain atlas.

## 2 Definitions and modeling hypothesis

Axons are complex structures composed of several subparts, as detailed in Gibson and Ma (2011). In this work, we suppose that the axons are divided into a long-range trunk, which can span all brain regions, and several local tufts, as shown in Fig. 1. The point connecting a tuft to the long-range trunk is called the *common ancestor* of the tuft. Also, for modeling consistency, the long-range trunk cannot contain any terminal. Terminals are always part of a tuft, so it is possible that a tuft actually contains only one terminal section. The long-range trunk is built so that it connects specific brain regions targeted by the axon. These brain regions are registered in a brain atlas that divides space into voxels and associates data with each of them (brain region ID, local orientation, etc.).

**Fig. 1:**
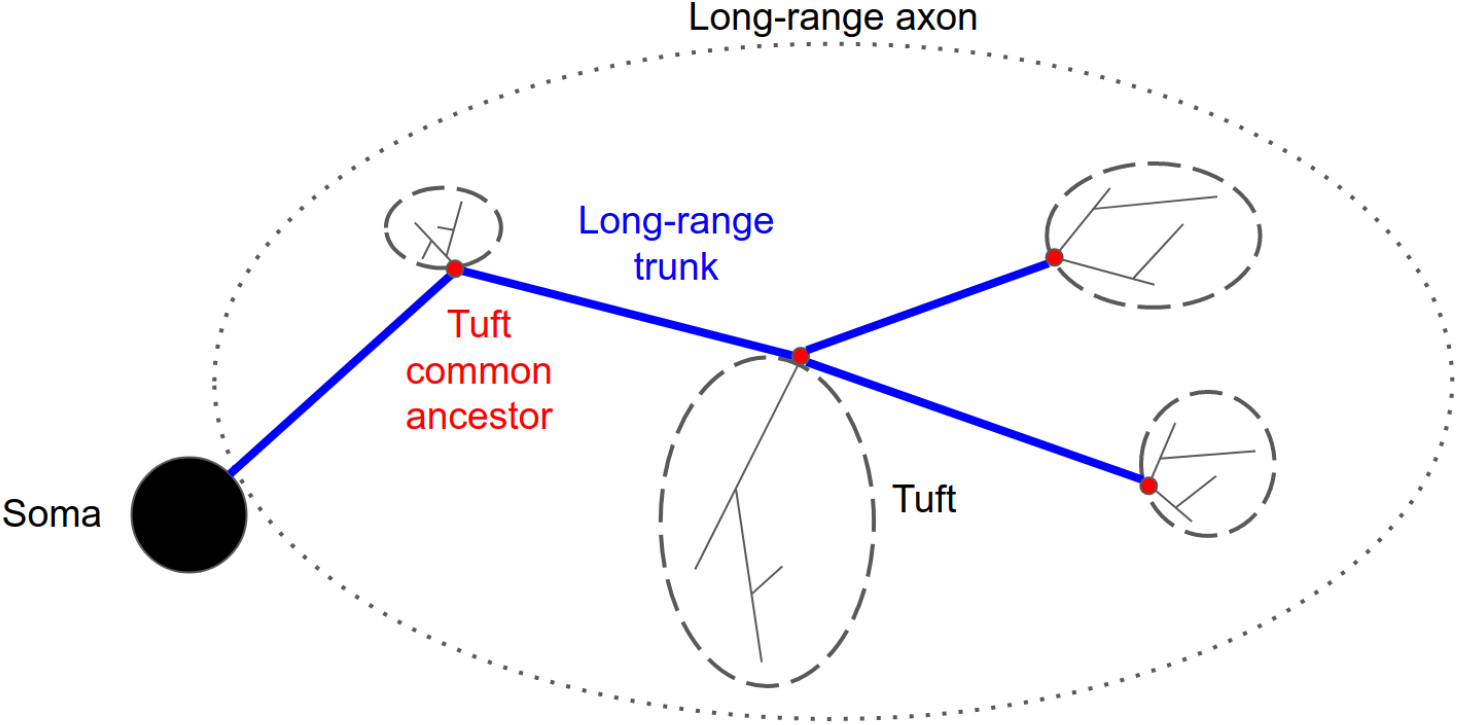
Schematic of long-range axon modeling, including the steps of the long-range trunk, the common ancestors, and the tufts generated at each target point.

This model is consistent with Gibson and Ma (2011) which states that during the development of the axon, a growth cone extends the axon in the direction of the targeted brain regions to connect to them. During this growth, branching processes occur in order to connect several brain regions while minimizing the total length of the axonal tree. Then, when the growth cone reaches a targeted brain region, the branch splits into multiple short terminal sections that will actually connect to the dendrites of the surrounding neurons. In this work, we aim at creating morphologies similar to the final state of the axon only, thus the intermediate states of the development are out of scope.

## 3 Methods

This section presents all the details of the method. The first subsection presents the data preprocessing used to calibrate the parameters required for the synthesis process. This synthesis process is described in the two following sub-sections. The first is dedicated to the synthesis of the main trunk and the second is dedicated to the synthesis of the tufts.

### 3.1 Creating statistical inputs from reconstructed cells

In order to be able to synthesize realistic morphologies, it is required to calibrate multiple parameters from real reconstructed neuron morphologies. There are two kinds of these parameters: statistics on local properties of the sections of both the long-range trunk and the tufts, and the global tuft properties that describe their sizes and shapes depending on where they are located.

First, in order to compute the statistics and properties of the long-range trunk and the tufts, it is required to define which parts of a reconstructed morphology are considered as tufts, and then the rest is considered as the long-range trunk. The tufts can be defined from multiple methods. In this work, they are defined using a simple clustering algorithm: two points separated by a Euclidean distance *d ≤ d*_max_ and a path distance 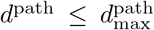 are considered part of the same tuft. Applying this algorithm to each pair of terminals and merging the pairs with a common ancestor defines all the tufts. Other clustering algorithms were tested (e.g., do not consider the path or define the tufts according to brain regions), but the one presented here was simple and gave good results.

Once the tufts are defined, it is possible to extract the required features from them. For each tuft, its *barcode*, which is a topological signature as defined by Kanari et al. (2020), is computed using the Topological Morphology Descriptor (Kanari et al., 2024). Then the following tuft properties are computed:

- the coordinates of the common ancestor,
- the total path length of the tuft,
- the path distance from the soma to the common ancestor,
- the orientation in the local atlas frame of the segment between the common ancestor and the center of mass of the tuft (the local atlas frame has the Y axis towards the pia).

All these properties are then stored in a dataset from which entries will be sampled to synthesize new tufts.

Finally, the mean and standard deviation of the lengths of the segments of the long-range trunk are extracted to make it statistically realistic during synthesis. These properties will be used to post-process the skeleton of the long-range trunk, as the main process described in 3.2.2 only aims at drawing its trajectory, not its actual morphology.

### 3.2 Building the long-range tree

The main trunk of an axon, also referred to as the long-range trunk or long-range tree, is the large-scale part of the axon that connects different brain regions. The synthesis process of the main trunk has 3 main steps:

1. find the source and target point coordinates;
2. connect the target points;
3. post-process the resulting tree.

These steps are presented in the subsections 3.2.1, 3.2.2 and 3.2.4.

#### 3.2.1 Source and target points placement

The long-range axon synthesis algorithm starts by collecting the given morphologies from the given cell collection to which the synthesized long-range axons should be grafted. Then the starting points of each axon are computed. These starting points are called the source points of the axons and can either be given explicitly (e.g. to start from an existing local axon) or just start from the soma.

The next step is to associate a *source population* to each long-range axon. These source populations represent groups of neurons that participate in projections to a given set of target brain regions. Source populations can be arbitrarily associated to each axon or computed using a brain atlas, such as Wang et al. (2020). In the latter case, the brain region of the atlas in which the cell is located is computed, then a source population is randomly picked among the ones associated to this brain region (as multiple source populations can be associated to a given brain region to represent the diversity of projection classes). The probabilities of these source populations are given in the input *source population matrix*, which can be derived from connectomics results or from large reconstructed morphology datasets (see Petkantchin et al. (2024) for an application of this work).

After that, the target populations of each axon are randomly picked among the ones possible for the associated source population, each target population possibly being associated with several brain regions. The probabilities of each target population depending on a source population are given in a *projection probability matrix*. Once the target populations have been chosen, a voxel is randomly chosen in the brain region associated with each target population. Then a random shift is applied inside this voxel and the resulting location is called a *target point* of the axon.

#### 3.2.2 Connecting source points to target points

As described by Cuntz et al., the Ramón y Cajal’s hypothesis about wiring optimization leads to near-optimal trees regarding the minimization of the total path length of the morphology. In graph theory, the problem consisting in connecting a set of terminal points (which are the target points described in 3.2.1) in Euclidean space while minimizing the total length of the graph is known as the Euclidean Steiner Tree Problem. This problem is known to be NP-hard, which is an issue to generate large axon structures in a reasonable amount of time. Also, in this problem, it is supposed that the cost function is uniform in space, which is not the case in our problem, as axons tend to follow axonal projection tracts for example. For these reasons, we use an approximation to connect the set of terminal points. As a first step, we transform the Euclidean problem into a graph problem by creating a 3D network connecting all the terminal points with some intermediate points. Then, the edge weights are manipulated to model different constraints (especially to decrease the weight of edges located in the projection tracts). Finally, the Steiner Tree is solved in this graph using the approximation described by Hegde et al..

The graph creation is divided into several parts. First, the terminal points are collected, and then several types of intermediate points are added. The first type consists in evenly splitting each segment between the source point and the target points into sub-segments to create *N*_intermediate_ new points. These splitting points are used to refine the future graph and to ensure it is possible to connect the terminal points using non-direct paths. Then *N*_random_ random points are added in the bounding box of the previous points so that there is no other point within a *r*_*random*_ radius. These random points are also used to refine the future graph and to ensure that the Steiner Tree algorithm can find the paths that appear longer in Euclidean space but have smaller weights because they go through preferred areas (e.g. projection tracts, see section 3.2.3). Then Voronoï points are computed between all the previous points and added to the set (the ones that are too far from the bounding box are discarded). These Voronoï points allow us to reduce the empty areas and ensure that the bifurcation angles are not too small, which is more realistic. Finally, close points are merged to reduce the size of the point set and to avoid too small segments. This process greatly increases the size of the set of points used to compute the Steiner Tree but gives a lot of possible paths to connect the target points and thus can give relevant solution paths.

Once the set of points is created, a triangulation is performed to create the edges of the graph. Then the edges of this graph are weighted according to Eq. 1, where the edge *i* connects the points *A* and *B, l*_*i*_ is the length of the edge *i, o*_*i*_ depends on the orientation of the edge *i, d*_*i*_ depends on the depths of the edge *i* in the atlas and *γ*_*i*_ depends on the space area in which the edge *i* is located. The first term of the weight is required since the total length should be as close to the minimum as possible to build a relevant tree. The second term is used to reduce the probability of transverse sections, which are less common. The third term is used to favor paths that follow the curvature of the atlas layers. The last term is required to favor specific areas (e.g. projection tracts).

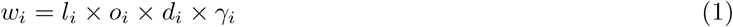

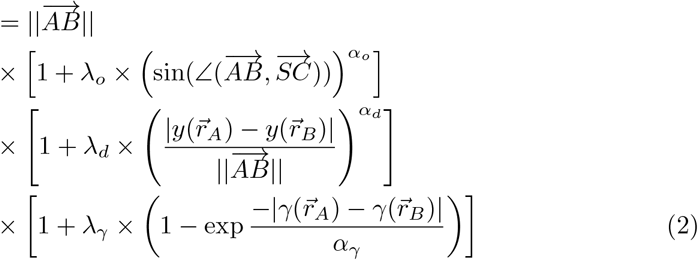

In Eq. 2, *S* is the center of the soma, *C* is the middle of the segment *AB*, 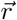 is a location in space, *α*_*o*_, *α*_*d*_ and *α*_*γ*_ are positive exponents, *λ*_*o*_, *λ*_*d*_ and *λ*_*γ*_ are amplitude coefficients, *y* is the depth field associating an atlas depth to each location 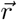 in space and *γ* is an attraction field that represents the affinity of the axons for each location 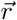 in space.

After these steps the Steiner Tree is computed on the resulting graph to select which edges are part of the solution and these edges are gathered to build the long-range trunk skeleton. Then, this skeleton will be refined to make small-scale morphometrics more realistic. A simple example of the long-range trunk skeleton building process is shown in Fig. C2a.

#### 3.2.3 Preferred region modeling

As described in Thiebaut de Schotten et al. (2011), the main trunks of the axons usually do not go straight to their targets. In contrast, they tend to use close trajectories in brain regions called axonal projection tracts. The Eq. 2 presented how the weights of each edge of the graph are assigned before computing the Steiner Tree. In this equation, the last term allows us to consider a set of attractors at arbitrary locations which generate a global attraction field. This attraction field can be used to guide the main trunks of the axons to make them pass through specific regions in the brain, as it is observed in projection tracts for example. Fig. C2b presents the same example as in Fig. C2a except that one attractor was added in the upper left corner. In this figure, edges are colored by the linear densities of their weights, where blue-colored edges have small weights and are thus preferred by the Steiner Tree algorithm, while the red-colored edges have high weights and are thus less likely to be selected by the Steiner Tree algorithm. As one can see, edges close to the attractor have smaller weights than edges far from the attractor. As a result, the optimal solution selected by the Steiner Tree algorithm, in this case, is very different from the one without any attractor: the first part of the main trunk passes close to the attractor and then goes to the closest target, and then it connects the furthest target. This simple example demonstrates that it is possible to alter the main trunk shape by defining attractors in space.

In the context of a synthesized axon, multiple attractors may be needed to represent the relevant attraction field. To test and validate the attraction mechanism, an axon was synthesized such that it should be very close to a given reconstructed axon, see Fig. C2c and Fig. C2d. In that case, the target points were defined as the common ancestor of each clustered tuft, as described in 4.1. The attractors were built using a new dummy brain region that was added to the atlas composed of all voxels intersected by the reconstructed axon. Then one attractor was created at each center of the voxels of this dummy region. When there is no attractor (Fig. C2c) the main axon trunk goes straight to the target, while when there are attractors (Fig. C2d) the trunk follows them properly. This example demonstrates that this method can follow complex paths in space when required data are available.

#### 3.2.4 Post-processing

The previous section 3.2.2 described how the long-range trunk skeleton is built to connect the proper brain regions. However, the resulting tree needs post-processing to ensure that its local morphometrics are statistically consistent with respect to reconstructed axons. This post-process consists of performing a correlated random walk guided by the intermediate points of the Steiner Tree. This random walk must be calibrated so that the result satisfies the local morphometrics extracted in 3.1.

An important feature for the subsequent growth process of the tufts is to keep immovable the initial target points that will be used as root points to grow the tufts (these root points will thus become the common ancestors of the synthesized tufts). For this reason, the steps described in this section all preserve these points in the morphology. Furthermore, the bifurcation points are also kept immovable for simplicity.

For each tree, a set of long-range trunk statistics is randomly selected among the ones extracted in 3.1. Then a guided correlated random walk is performed between each consecutive immovable point in the tree, whether they are the source point, a bifurcation point, or a target point. The inputs of this random walk are thus a start point, an end point which is the global target to reach, and a set of intermediate points which are intermediate targets close to which the random walk must pass. The direction of each step of this random walk is defined by Eq. 3, where 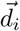 is the unit vector pointing in the direction of the current step *i* and which is equal to a weighted sum of several unit vectors: 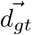 is the unit vector pointing to the endpoint, 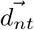 is the unit vector pointing to the next intermediate point, 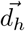 is computed from the history of the previous steps and 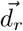 is a random unit vector. The *β*_*i,gt*_, *β*_*i,nt*_, *β*_*i,h*_ and *β*_*i,r*_ are the weights applied to each associated term. These weights depend on the current step.

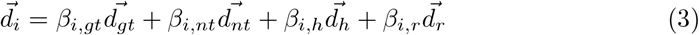

At each step, the distance to the next intermediate target is evaluated and if it increased since the previous step or if it is smaller than the mean step length multiplied by a given coefficient (equal to 1 in this work), then the intermediate target is considered as reached and the next one becomes the new intermediate target. This process ensures that the random walk follows the long-range trunk skeleton as defined in 3.2.2.

After these post-processing steps, the long-range trunk is statistically close to the reconstructed ones and is thus ready to be used to grow the tufts from it.

#### 3.3 Tufts

This section presents the method used to create the tufts and graft them to the long-range trunk. For a given tuft, the general process consists of synthesizing a new tuft based on the properties of a reconstructed tuft. This reconstructed tuft is chosen as a template from a set of tufts according to a given weight. This weight is computed for each target population, as defined in 3.2.1, from a set of properties that should be as close as possible to what is needed for this newly synthesized tuft. These properties can be based on the morphometrics of the tuft or contextual properties. For example, simple morphometrics can be the path length from the soma (*d*_*i*_) or the total path length of the tuft. A contextual property can be the brain region in which the tuft is located or the distance of the common ancestor of the tuft from the pia.

For example, using the path distance from the soma to the initial point and the expected total path length of the tuft, a probability can be calculated for each reconstructed tuft that may be chosen as a template. For each potential template *j*, an affinity is computed as:

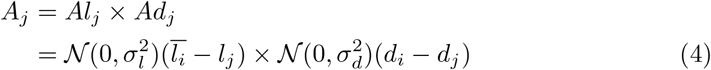

where *A*_*j*_ is the affinity of the template *j, Al*_*j*_ is a component of the affinity associated with its total path length, *Ad*_*j*_ is the affinity associated with its distance from the soma, *N* is a normal distribution, *σ*_*l*_ is the given standard deviation for the expected total path length of the tuft and *σ*_*d*_ is the given standard deviation for the path distance from the soma. Then the probability *P*_*j*_ to choose a template *j* is computed as follows:

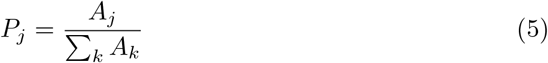

Once a template tuft is chosen, a new tuft is synthesized using the <monospace>NeuroTS</monospace> algorithm Kanari et al. (2022). The barcode given as input is the one of the template tuft, so the total length of the synthesized tuft is expected to be close to the one needed. In this work, all synthesized tufts use the same set of parameters and distributions in the <monospace>NeuroTS</monospace> algorithm, but it is possible to use specific ones for each tuft to refine the results.

Finally, the synthesized tuft is grafted onto the target point which becomes its common ancestor.

The main steps described in this section, from the target definition to the generation of tufts, are summarized in Fig. 2.

**Fig. 2:**
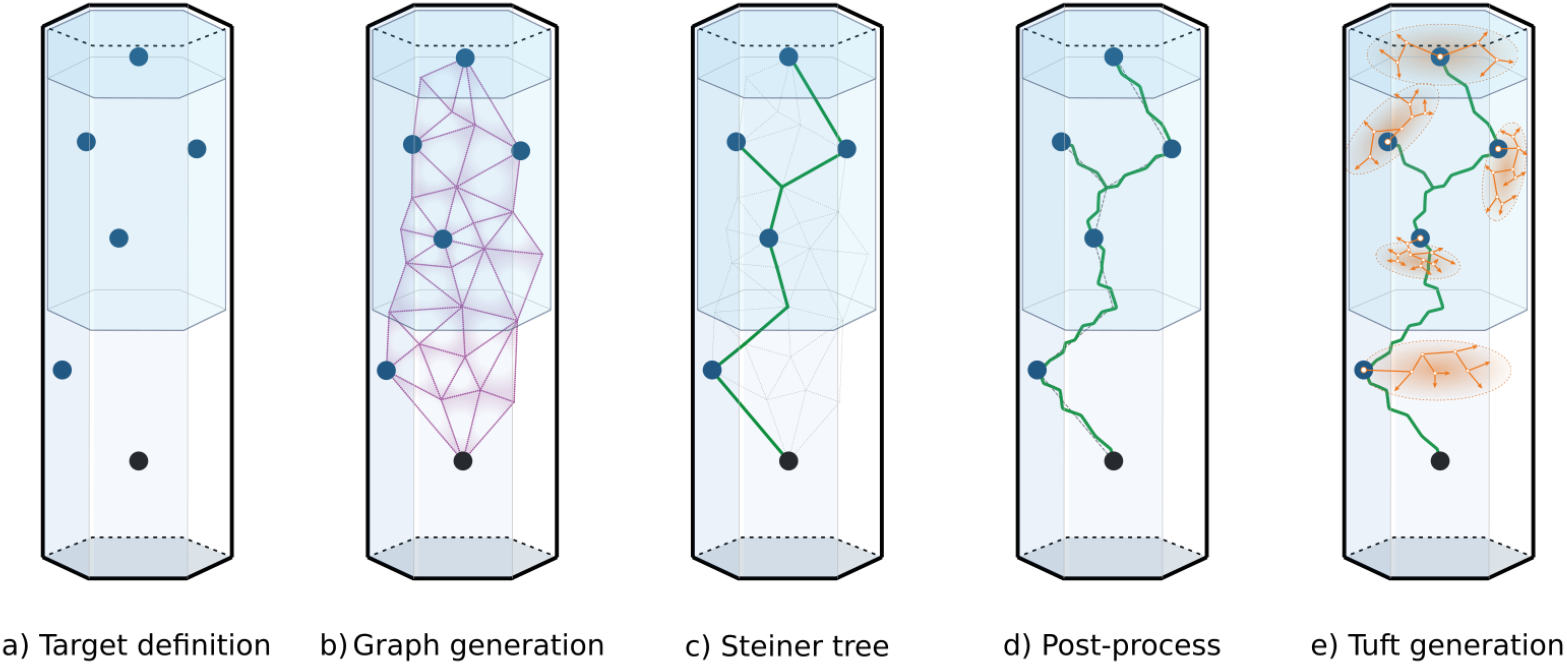
Main steps of the long-range axon synthesis algorithm: define the target points (a), generate a 3D graph containing the source point, the target points and other intermediate points (b), compute the Steiner Tree that connects the source point to the target points through this network (c), post-process the Steiner Tree solution to make it more realistic (d) and generate one tuft at each target point (e).

## 4 Results

This section presents a validation process that demonstrates that it is possible to synthesize an axon that properly mimics a reconstructed axon. We then study the effect of an important input parameter: the clustering size. Finally, we show an example application consisting of using a set of reconstructed morphologies to compute the target points of the axons and using the atlas data to follow the projection tracts in a mouse brain.

### 4.1 Mimicking biological reconstructions

In order to verify that the method presented in Section 3 can build realistic long-range axons, the capacity to synthesize axons similar to reconstructed ones is evaluated.

The main steps of the so-called *mimic* algorithm, when used to mimic a reconstructed morphology, are summarized in Fig. 3. The first step, for a given reconstructed axon (Fig. 3a), consists in clustering its tufts and reducing them to the common ancestors of their clusters, as described in 3.1. Fig. 3b shows the biological reconstruction after the tuft clustering: tuft sections are represented in orange while the trunk is represented in green. Then, a graph is created as described in 3.2.2. In this case, the preferred regions as described in 3.2.3 are enabled. These preferred regions are constructed along the reconstructed morphology, as presented in 3.2.3. Then we use the Steiner Tree algorithm (3.2.2) to build the long-range trunk that will use the common ancestors of the clustered tufts as target points. This synthesized trunk is then post-processed as described in 3.2.4. Fig. 3c presents the synthesized trunk after this post-process. Finally, the tufts are synthesized as described in 3.3, based on the properties of the clustered tufts of the reconstructed axon. Fig. 3d shows the final result of the synthesis process.

**Fig. 3:**
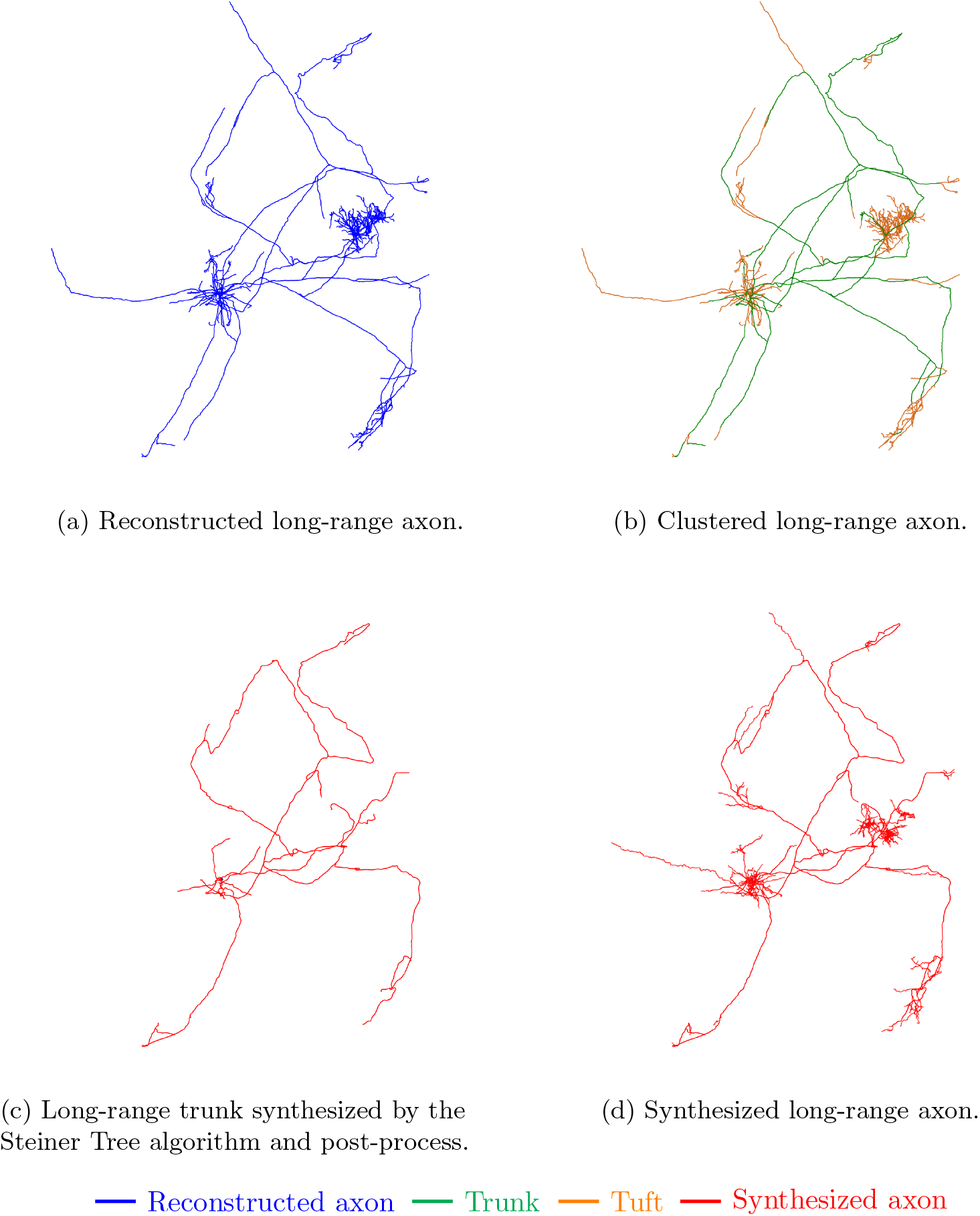
Main steps of the long-range axon mimic algorithm, including the steps of clustering the input morphologies (3.1), synthesizing the main trunk (3.2) and synthesizing the tufts (3.3).

These figures show that the global shape of the synthesized long-range axon is very close to the one of the initial reconstructed morphology. The long-range trunk goes to the proper target areas, and the tufts have similar sizes and orientations, so the target regions are properly innervated. Nevertheless, the branches of the long-range trunk are not identical. This is expected because the intermediate points given to the Steiner Tree algorithm are not only based on biological features and are partially random, so the result can not be the same as the reconstructed morphology. Also, one can see that the parallel trunks seen in 3a (e.g., in the lower-left part of the morphology) are replaced by a single trunk in 3d, this is because the Steiner Tree Algorithm tries to minimize the total length of the trunk, so the parallel trunks can only appear if there is a forbidden area between them. This very specific feature is not covered in this work as it is hard to get such data and because we suppose that the trunks have a small contribution to the connection network compared to the tufts.

In addition, Fig. B1 shows that most local morphometrics are consistent between the synthesized and reconstructed axons. Some morphometrics are not consistent, especially the mean section lengths and, to a lesser extent, the inter-segment angles. The section length is expected to be different because the bifurcations in the trunk have no reason to be located at the same places.

Fig. 4 shows several examples of synthesized axons that mimic reconstructed ones. The clustering distance is 100 *μ*m for all of them. Again, the results show good accuracy.

**Fig. 4:**
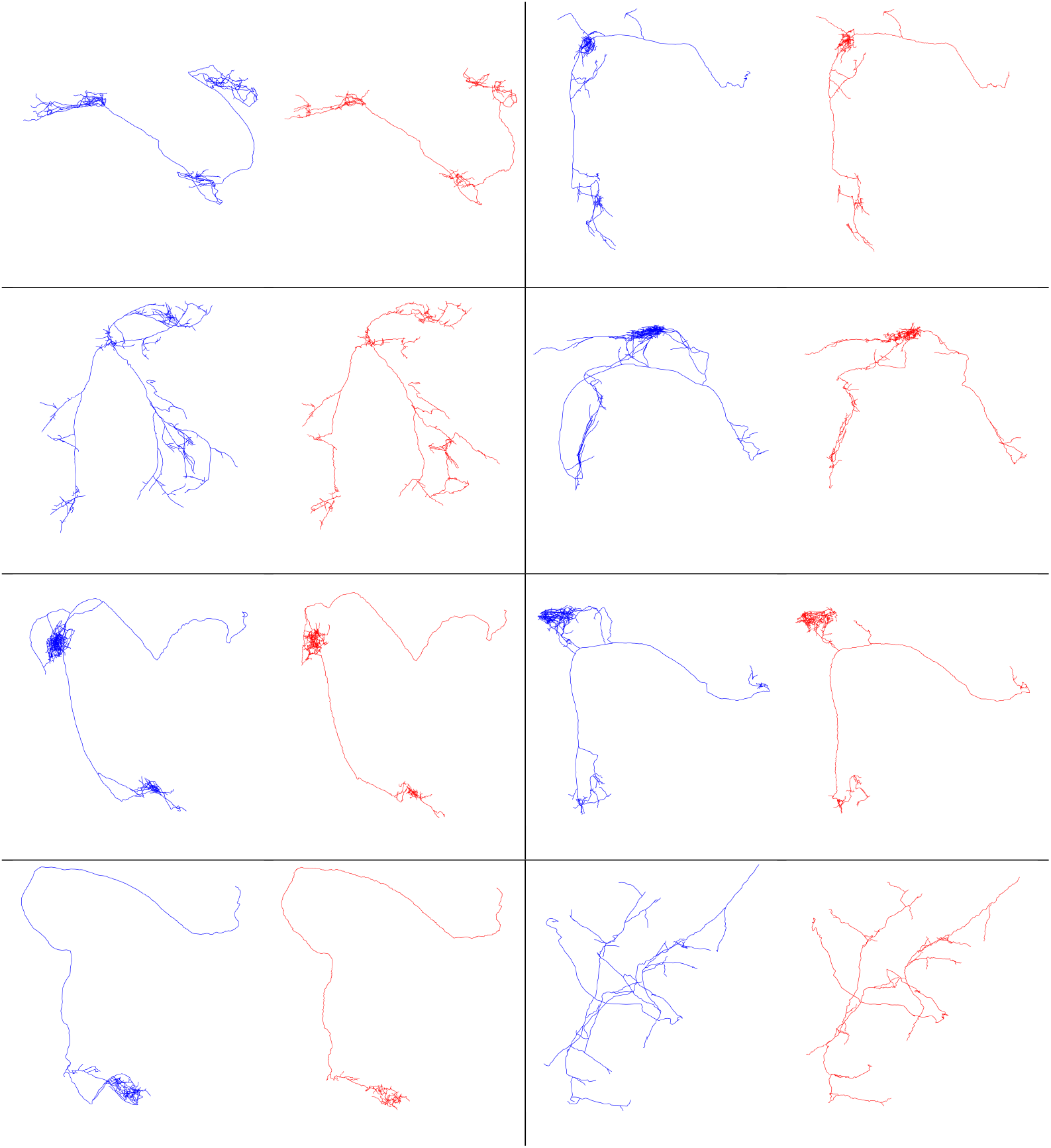
Examples of synthesized long-range axons (red) and the respective reconstructed axons (blue). These results demonstrate the ability of the algorithm to approximate a variety of different shapes.

In addition, to verify that the synthesized morphologies are statistically similar to the biological reconstructions, we mimicked 1076 morphologies of the MouseLight project (Winnubst et al., 2019). These morphologies, extracted from adult mice (age > 8 weeks), originate and target various brain regions. We compared the distributions of morphometrical features across the reconstructed and synthesized populations. These distributions for the trunks, tufts, and complete morphologies are plotted in Fig. 5. The morphometrics used are defined in the Petilla terminology (Ascoli et al., 2008; Kanari et al., 2022). We also plotted the Maximum Visible Spread (MVS) score, defined in Kanari et al. (2022), for the trunks, tufts, and complete morphologies in Fig. 5. This score gives an indication of the similarity between two statistical distributions, by computing the absolute difference between the medians divided by an estimate of the overall visible spread of the distributions. This score is minimal at 0 when the medians coincide and the distributions are very close to each other; the closer it gets to 1, the less similar the distributions are. The morphometrics of the reconstructed and synthesized axons match very closely at a population level, with the worst MVS of 0.12, 0.20, and 0.16 for the bifurcation angles of the trunks, tufts, and the final morphologies, respectively.

**Fig. 5:**
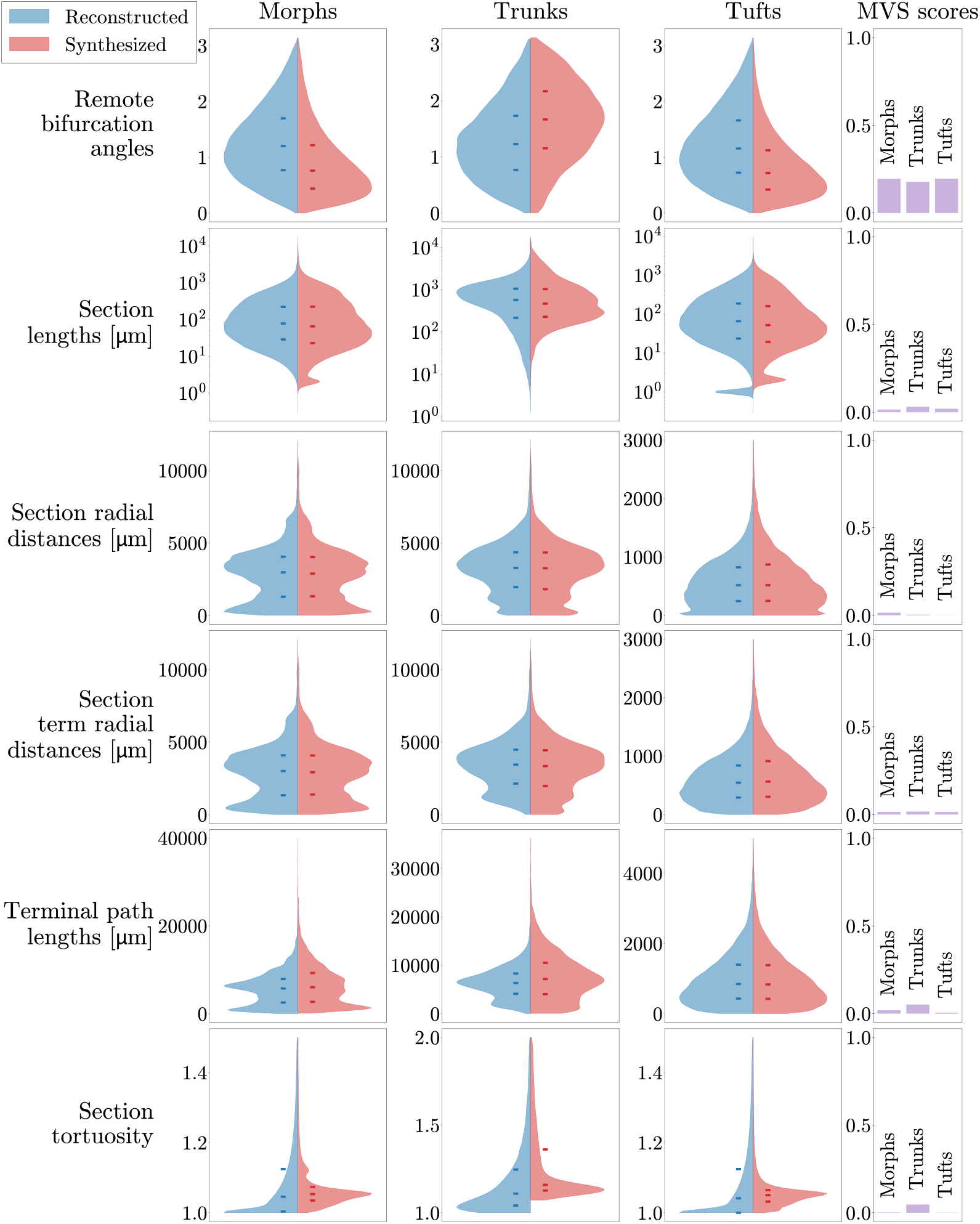
Distribution of morphological features across 1076 reconstructed (blue) and synthesized (red) axonal complete morphologies (left column), trunks (middle column) and tufts (right column). Median, first and third quartiles are shown on the distributions (hyphens). The MVS statistical score is shown in the last column for each type of morphology.

### 4.2 Clustering distance effect

The clustering distance used to define the tufts in the reconstructed morphologies is a crucial parameter when aiming to achieve results that closely match the reconstructed morphology. Small clustering distances tend to yield more detailed and accurate representations. However, in practical applications, using such small clustering distances can be challenging due to data accuracy and depends on how the target points are defined. As a result, it becomes necessary to find a compromise between the desired accuracy and the practical feasibility. Fig. 6 shows the effect of the clustering distance on a given synthesized axon compared to the corresponding reconstructed axon. As can be seen, the synthesized axon is very close to the reconstructed one when the clustering distance is small (6a) while it becomes less and less accurate with higher clustering values (6b to 6d). Fig. D3 shows the normalized *L*1 error of the projection intensity for each clustering distance. The projection intensity plotted in this figure is computed as follows: a 3*D* grid is created with a given voxel size, then the intersection of the axon and each voxel is computed for both the reconstructed and the synthesized axons. Then the normalized *L*1 error between the voxels of the two previous results is computed with Eq. 6.

**Fig. 6:**
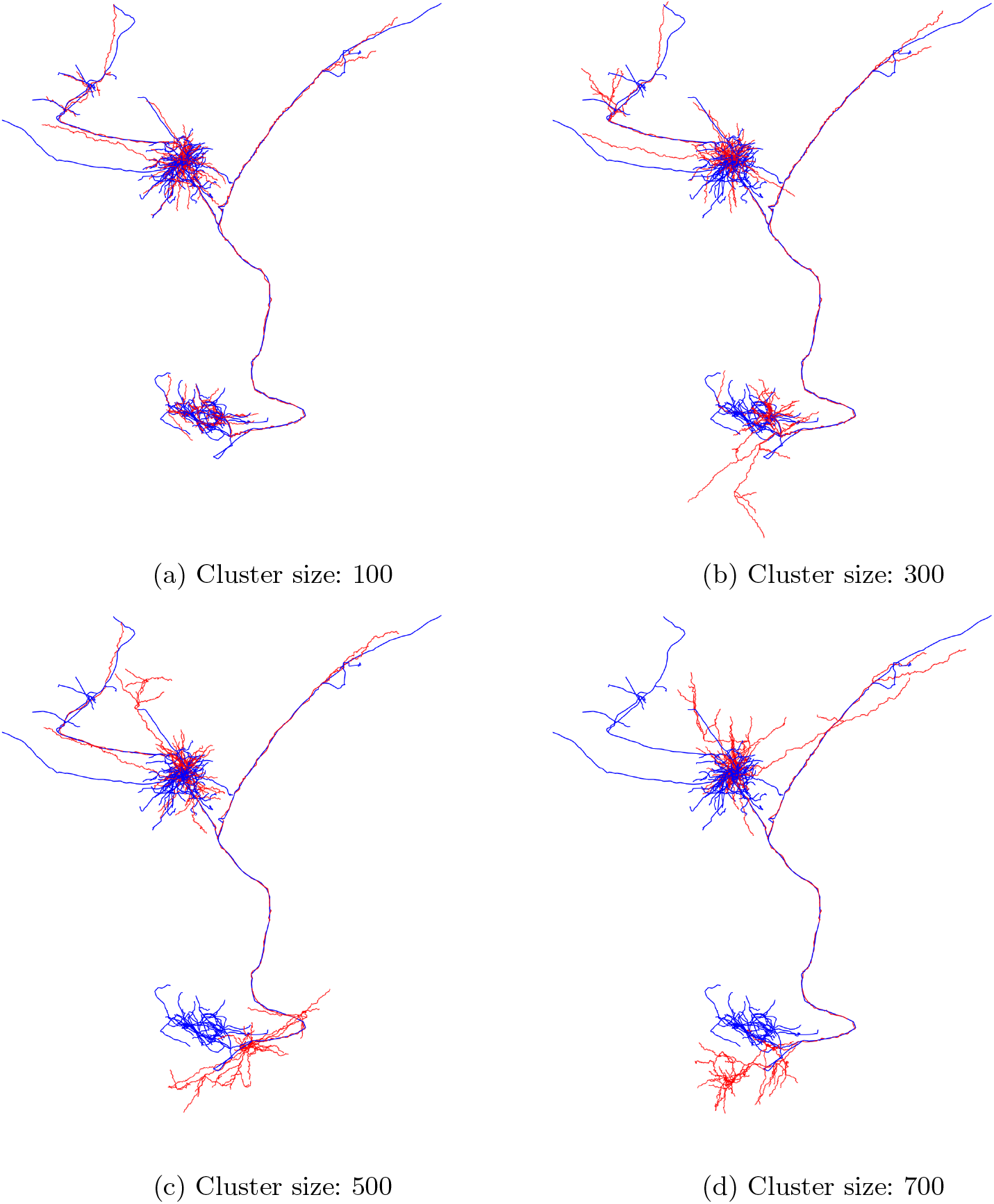
Parameter selection for cluster size. Synthesized long-range axons (red) for varying cluster sizes compared to reconstructed axons (blue). Smaller cluster size approximates the original axons with higher accuracy.

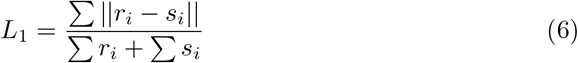

The process is repeated for multiple voxel sizes in order to show which scales are properly synthesized and which are not. The result shows that for a clustering distance equal to 100 *μ*m the synthesized axon has very similar intersection lengths as the reconstructed axon as long as the voxel size is bigger than 250 *μ*m, while the error increases fast for low voxel sizes. This means that the spatial structures bigger than this scale are accurately synthesized while the smaller structures are less accurate, which is expected since the synthesized axon is only supposed to be similar to the reconstructed one but not strictly equal. For bigger cluster sizes, the size of accurately synthesized structures is much bigger.

### 4.3 Application: connecting mouse brain regions

All previous tests have shown that the presented model is capable of accurately synthesizing long-range axons in many different situations. This last section presents a possible use case in which reconstructed morphologies are used to define the target point locations in the brain atlas. In this example, 44 morphologies from Winnubst et al. (2019) with somata located in the layer 5 and 6 of the Primary Motor Area (MOp5 and MOp6 regions) were clustered to define trunks and tufts. Random points were then selected around these tuft locations using a normal distribution with a standard deviation equal to 200 *μ*m, while ensuring that the selected points remained in the same brain region as the initial tuft point. The properties of the tufts were directly used for picking the barcodes during the synthesis of the new tufts.

#### 4.3.1 Projection tracts as preferred regions

Using this set-up, we first checked the effect of the preferred region feature. Here, all the brain regions considered as projection tracts in the CCFv3 atlas (Wang et al., 2020) are used as preferred regions. In this context, two long-range axons were synthesized using the same source and target points, the only difference being whether the preferred regions were enabled or not. As can be seen in Fig. 7, the two axons are very different: when the preferred regions are disabled, the main trunk goes straight to the target points, while when they are enabled the trunk follows the projection tracts when possible and then goes to the target points. This example shows that the model is able to follow realistic projection tracts in the brain, which is a key feature for realistic long-range axon synthesis.

**Fig. 7:**
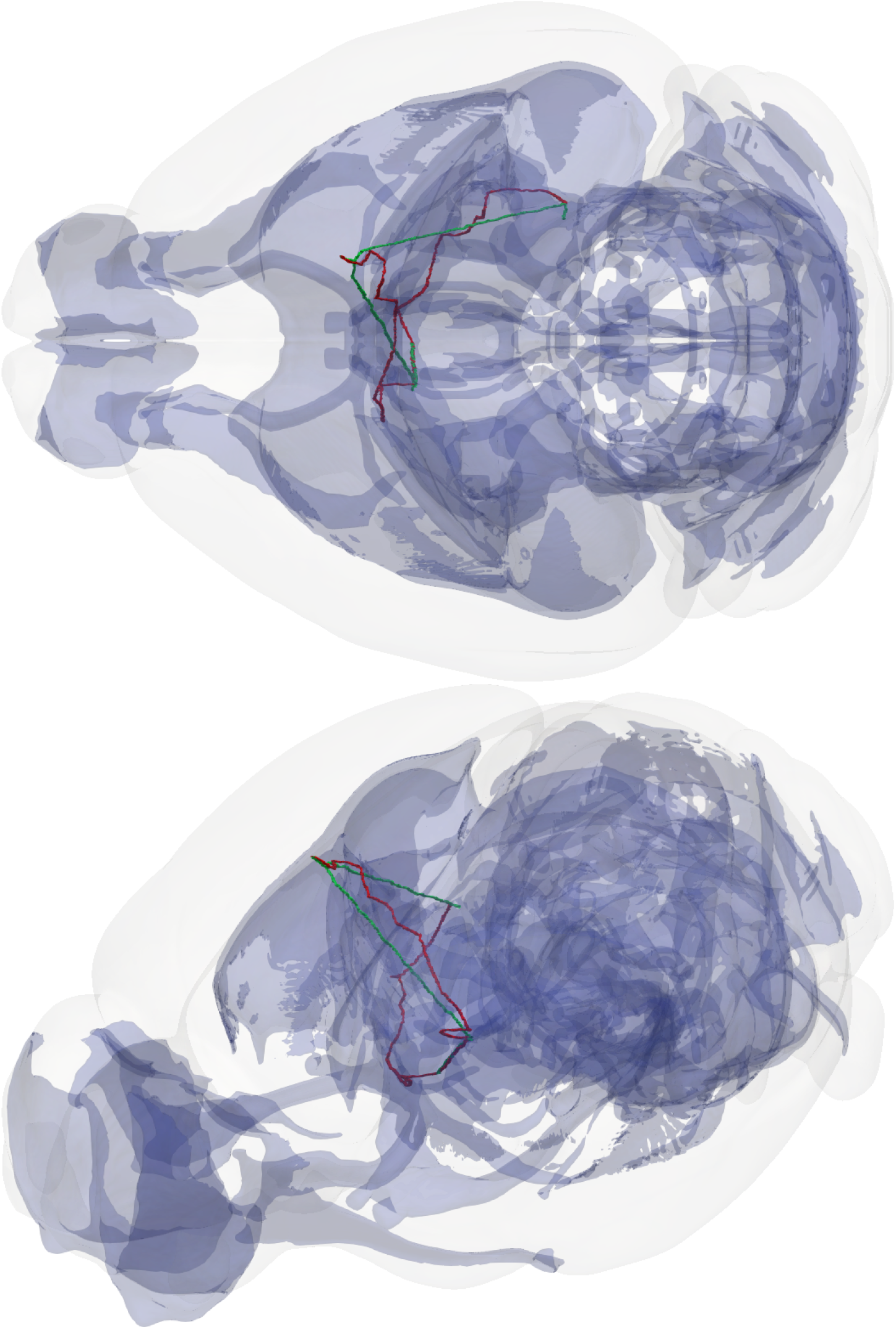
Comparison of two synthesized long-range axons, whose soma is located in the Primary Motor area, within the brain Atlas. The preferred region feature is disabled for the green one while it is enabled on the projection tracts for the red one.

#### 4.3.2 Boundary constraints

This application aims at synthesizing long-range axons in a specific spatial context, namely that of a mouse brain. In this context, an important feature is to ensure that the synthesized morphologies do not go outside of the brain. In this work, this is done by applying simple boundary constraints. The first constraint only applies to the long-range trunk since it is applied during the graph creation step described in 3.2.2. During this step, all the points of the graph located outside the brain are discarded before computing the Steiner Tree. This ensures that the long-range trunk does not go outside the brain. The second constraint only applies to the tufts, since it is applied during the tuft generation step described in 3.3. First, the space is divided into a 3D grid, the same as the one containing the atlas brain regions. Then for each voxel of this grid, the vector from the center of the voxel to the closest boundary point is calculated, so we obtain a 3D vector field, called the *boundary field*, each vector representing the distance and direction to the closest boundary point. Finally, the vectors located outside the brain are reversed and resized to a small length (1*e −* 3 *μm* in this work) to make the space outside the brain repulsive. After this process, the growth of the tufts is modified so that at each step of the random walk the distance and direction of the current point to the closest boundary are extracted from the previous boundary field. This distance is then refined to consider the position inside the voxel and used to calculate an attenuation vector using the following equation:

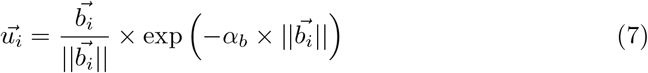

where 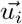 is the attenuation vector for the point *i*, 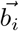 is the vector from the boundary field at the location of the current point *i* and *α*_*b*_ is a scaling coefficient. The refined distance is calculated by a simple linear extrapolation, considering that the attenuation field keeps the same direction inside a voxel. So, the vector from the current point to the center of the voxel can just be projected on the vector associated with the voxel and added to it to refine the distance to the boundary.

Finally, this attenuation vector is added to the direction of the current step of the random walk to reduce the component in the direction of the boundary. This process will reduce this component more and more so that it cannot cross the boundary.

#### 4.3.3 Multiple axon synthesis

Using the same setup, 44 long-range axons were synthesized. The model is again configured to follow the projection tract regions of the CCFv3 atlas (Wang et al., 2020). The main parameters given to the model are presented in Table A1. Fig. 8 shows that the long-range axons follow the projection tracts and properly innervate the same brain regions as the reconstructed morphologies. In this result, one can notice that the synthesized axons do not always follow the same tracts as the reconstructed morphologies. This is because all the tracts were considered equally attractive in this application for simplicity, but this is not the case in reality and should be refined for a real application.

**Fig. 8:**
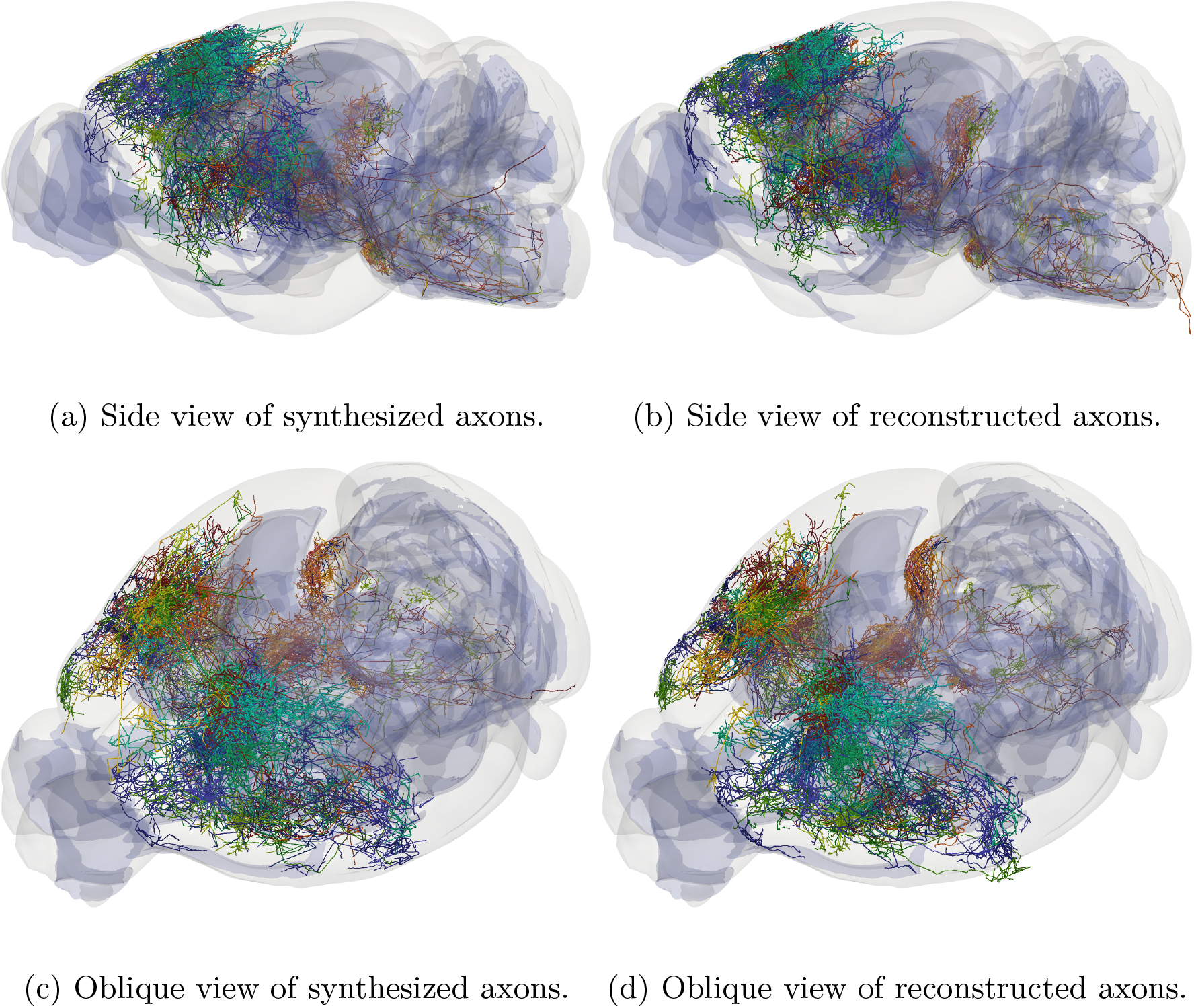
Example of 44 long-range axons synthesized such that follow the projection tracts (blue region) (a and c) compared to the respective 44 reconstructed morphologies (b and d).

Finally, we compared a set of morphometrics when boundary constraints were enabled or disabled to ensure that this process did not have a sensitive impact on the local structure of the morphologies. The Table 1 presents the MVS scores for several morphometrics of the morphologies synthesized without the constraint compared to those calculated with the constraint. As can be seen, most of these morphometrics are very similar (MVS scores are all very close to 0). Note that tortuosity and radial distances, are expected to be different since the sections close to the boundary are curved when the boundary constraints are enabled.

**Table 1:** MVS scores of the morphometrics distributions computed on the morphologies synthesized with and without the boundary constraints enabled.

| Morphometric name | MVS score |
| --- | --- |
| Section lengths | 0.000772 |
| Remote bifurcation angles | 0.009667 |
| Section radial distances | 0.002622 |
| Section term radial distances | 0.002159 |
| Section branch orders | 0.000000 |
| Section tortuosity | 0.001269 |
| Terminal path lengths | 0.004756 |

### 4.4 Benchmarking

The method presented in the previous sections was developed to be computationally efficient. To estimate its computational speed, a test was performed in which 100000 axons were synthesized on a cluster with 200 tasks. The morphologies started from the Primary Motor Area (MOp) and expanded to multiple other regions in the brain. parameters were consistent with those used in 4.3.3. In these conditions, the test is completed in 33min 15s. The total length synthesized considering all axons was equal to 1.01*e*3m, which means that the mean length of the synthesized axons is equal to 1.01*e*4*μ*m. This computational speed enables one to synthesize large neural circuits in a reasonable time.

## 5 Discussion

The synthesis of long-range axons is a complex yet essential step in reconstructing complete brain networks. In this work, we introduce a novel algorithm to synthesize long-range axons accurately. The synthesized axons show strong statistical agreement with expected biological properties, including local morphometrics, global targeting, and intermediate trajectories through preferred regions, such as projection tracts. Thus, the presented algorithm enables the generation of a diverse range of axons contributing to the reconstruction of large-scale brain circuits. In addition, the method is computationally efficient, and can thus generate 100000 axons of approximately 1km total length in about half an hour on a cluster with 200 tasks.

The axon synthesis algorithm can be further refined to generate more accurate axonal morphologies and, importantly, to increase variability within computational models of long-range axons. The current version shows sensitivity to the quality of input data, as shown in 4.2. As more experimental data becomes available, the algorithm should generalize to capture the global properties of long-range axons to minimize dependency on specific experimental inputs. Additionally, the selection of projection tracts should be guided by the source and target locations of each axon, as outlined in Budd et al. (2010) and Thiebaut de Schotten et al. (2011), while also accounting for the different neuronal classes, as described in Liu et al. (2024). We expect that these refinements will not impact the computational cost of the methodology.

Another potential improvement is to modify the Steiner tree algorithm to achieve a balance between cable cost and distance from the source, following an approach similar to the Tree algorithm, as described in Cuntz et al. (2010). Along with optimizing the selection of Steiner parameters, we will also implement an enhanced version of source-target connectivity. These modifications will improve specific morphometrics that currently do not perfectly match experimental data, such as inter-segment angles, bifurcation angles, and branch orders.

In a companion study, Petkantchin et al. (2024), we focus on generalizing the axon targeting from the available biological inputs for each brain region and hemisphere, and adding specialized tuft selection and placement to avoid replicating individual morphologies, thereby more accurately capturing population variability. By adding a clustering process to better differentiate the synthesis inputs, we also capture the source and targets of axons more accurately, improving in addition the input positions to the Steiner Tree algorithm. These improvements, alongside the incorporation of increasing experimental data, will further enhance the quality and biological relevance of the synthesized axons. Additionally, this work facilitates the generation of detailed connectivity between brain regions, enabling the computational reconstruction of biologically accurate synthesized neural networks.

The proposed algorithm enables the generation of large-scale neuronal circuits with biologically realistic connectomes, facilitating extensive simulations on virtual brain models.

The axon synthesis algorithm enables the generation of large-scale neuronal circuits with biologically realistic connectomes, generating detailed computational brain models. This capability allows for in-silico experiments that are otherwise unfeasible in-vivo, providing a powerful tool for advancing neuroscience research. The integration of synthesized brain regions with electrical models further enables the study of signal propagation across large neural circuits. This step is essential for simulating brain disorders and investigating specific neurological conditions such as autism, epilepsy, Alzheimer’s disease, and other neurodegenerative disorders (Belmonte and Bourgeron, 2006; Kaufmann and Moser, 2000; Moolman et al., 2004; Srivastava et al., 2012; Torben-Nielsen and Cuntz, 2014). While the full validation of such models remains a challenge, this approach holds significant promise for improving early-stage diagnostics and accelerating drug discovery. Moreover, this translational approach offers the potential for personalized medicine, enabling patient-specific drug testing and therapeutic strategies over the long term.

## Appendix A Synthesis parameters

**Table A1:** Input parameters used to generate the long-range axons presented in the different sections of this work.

| Synthesis case | Synthesis step | Parameter | Value |
| --- | --- | --- | --- |
| Long-range axons presented in 4.3.1 | Clustering | $d_{min}$ | 200 |
| | | $d_{max}^{path}$ | 300 |
| | Graph creation | $\alpha_o$ | 0.1 |
| | | $r_{random}$ | 500 |
| | | $\lambda_o$ | 1 |
| | | $\alpha_d$ | 0.25 |
| | | $\lambda_d$ | 2 |
| | | $\alpha_\gamma$ | 10 |
| | | $\lambda_\gamma$ | 5 |
| Long-range axons presented in 4.3.3 | Clustering | $d_{min}$ | 200 |
| | | $d_{max}^{path}$ | 300 |
| | Graph creation | $\alpha_o$ | 2 |
| | | $r_{random}$ | 500 |
| | | $\lambda_o$ | 2 |
| | | $\alpha_d$ | 0.25 |
| | | $\lambda_d$ | 2 |
| | | $\alpha_\gamma$ | 30 |
| | | $\lambda_\gamma$ | 7 |
| Morphometrics presented in Fig. 5 | Clustering | $d_{min}$ | 300 |
| | | $d_{max}^{path}$ | 1000 |
| | Graph creation | $\alpha_o$ | 0.1 |
| | | $r_{random}$ | 500 |
| | | $\lambda_o$ | 1 |
| | | $\alpha_d$ | 0.25 |
| | | $\lambda_d$ | 2 |
| | | $\alpha_\gamma$ | 100 |
| | | $\lambda_\gamma$ | 1 |

## Appendix B Morphometrics

**Fig. B1:**
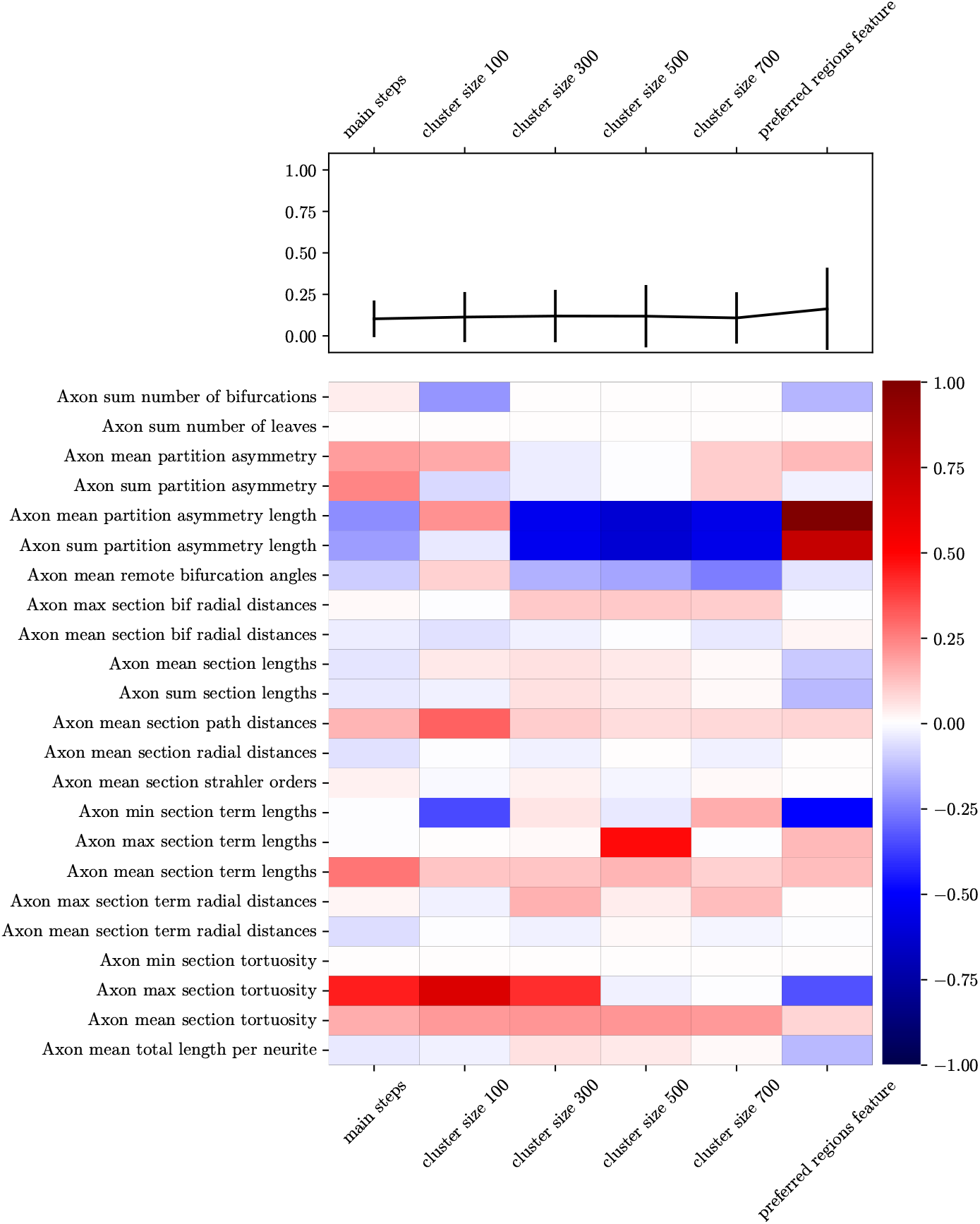
Errors on local morphometrics for different axons presented in this work. White cells mean that the morphometric values for synthesized and reconstructed axons are very close while red/blue cells mean that the synthesized value is much higher/lower. The upper part of the figure shows the global error on all the morphometrics with error bars. See the NeuroM documentation for details about morphometrics.

## Appendix C Graph creation and preferred regions

**Fig. C2:**
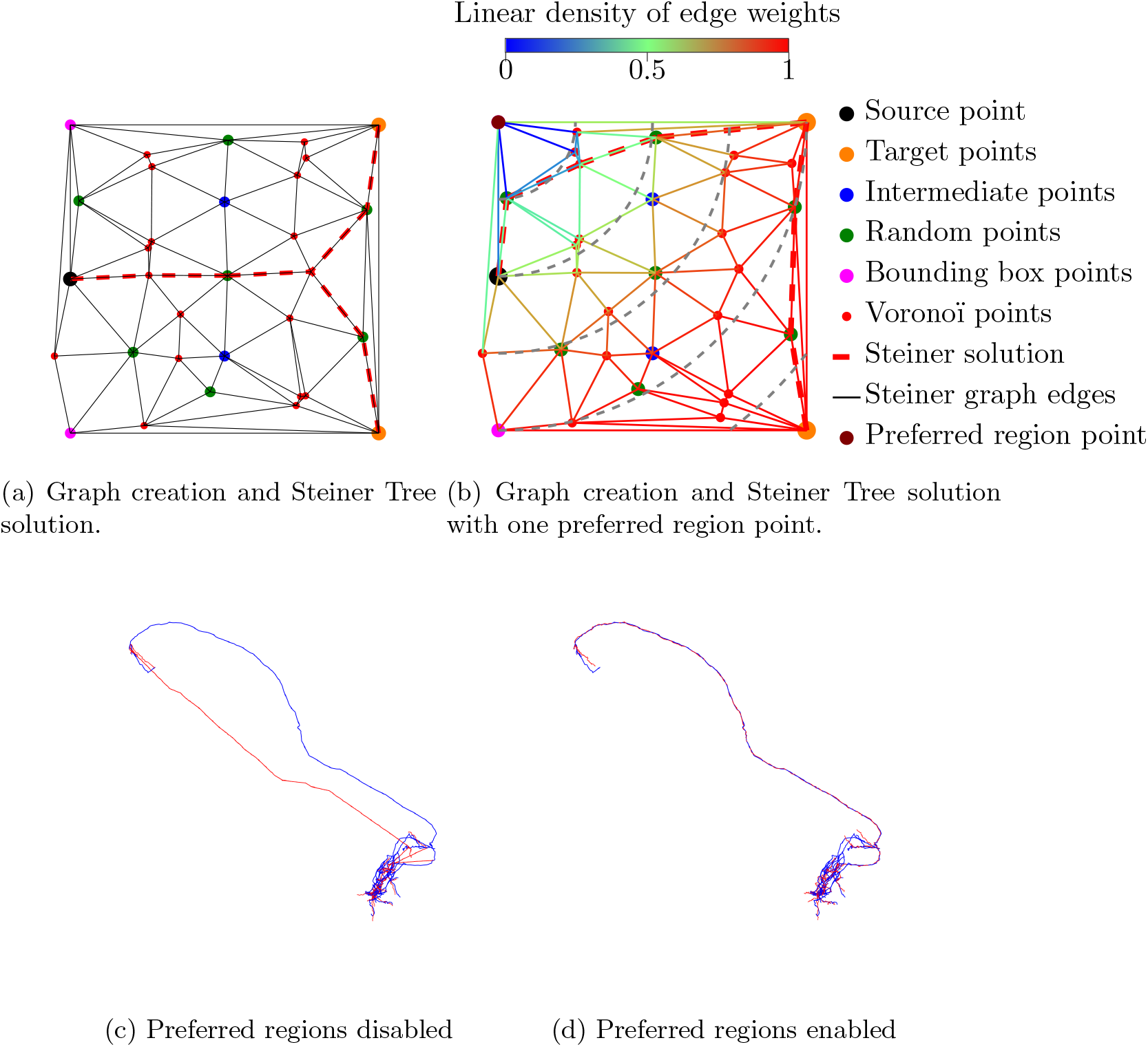
Example of graph creation on a simple case when there is no preferred region attractor (a) and with one attractor (b). The resulting synthesized long-range axons (red) compared to reconstructed axons (blue) are presented when the preferred regions are disabled (c) and enabled (d).

## Appendix D Error with cluster size

**Fig. D3:**
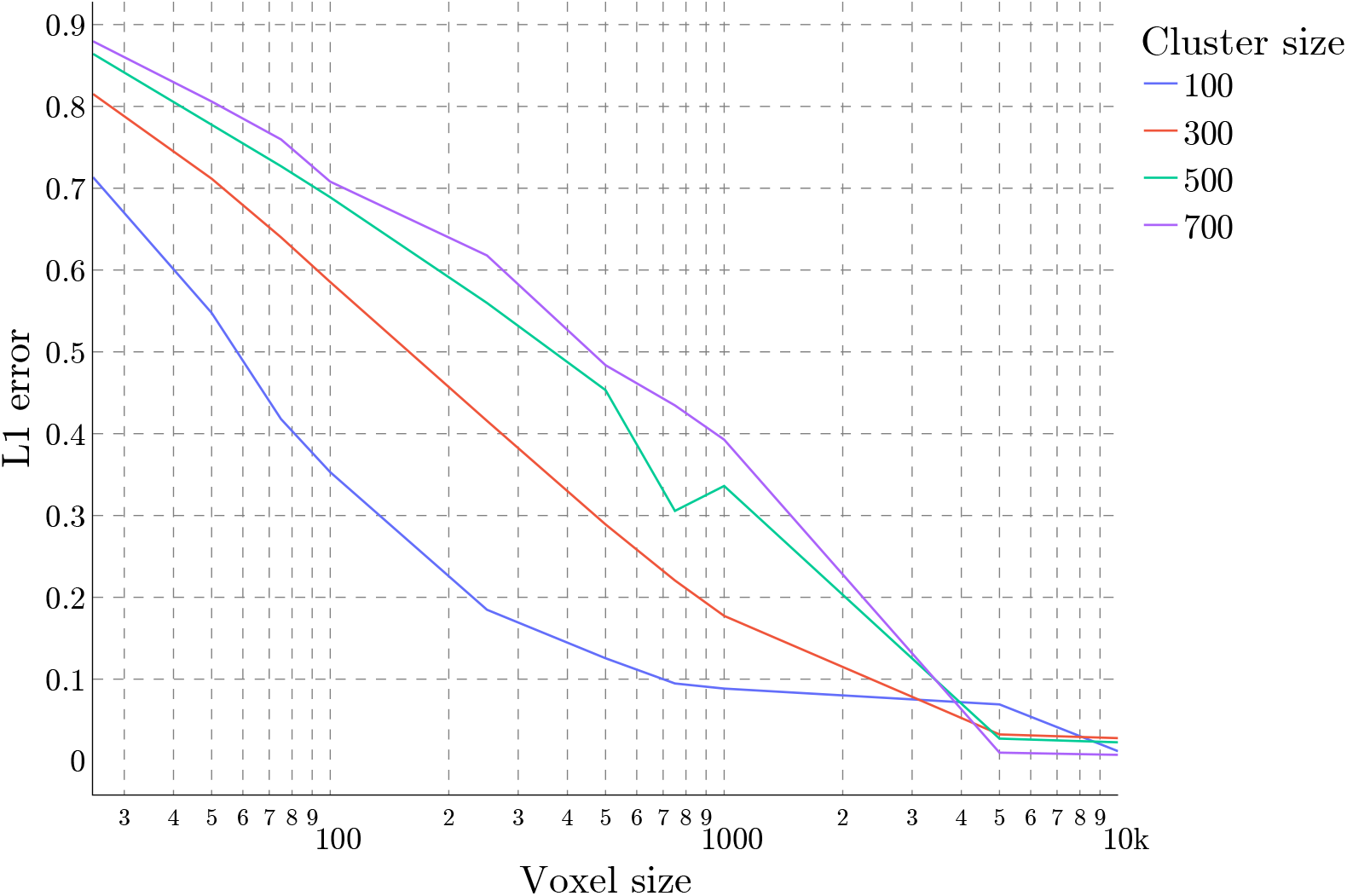
Projection intensity errors for different cluster sizes.

## Information Sharing Statement

All data and scripts used to generate the results presented in this work are available here: https://doi.org/10.5281/zenodo.13843538

The main code used for axon synthesis will be available here: https://github.com/BlueBrain

## Acknowledgements

This study was supported by funding to the Blue Brain Project, a research center of the École polytechnique fédérale de Lausanne (EPFL), from the Swiss government’s ETH Board of the Swiss Federal Institutes of Technology.

